# Reactive Oxygen Species Regulate Activity-Dependent Neuronal Structural Plasticity in *Drosophila*

**DOI:** 10.1101/081968

**Authors:** Matthew C. W. Oswald, Paul S. Brooks, Maarten F. Zwart, Amrita Mukherjee, Ryan J. H. West, Khomgrit Morarach, Sean T. Sweeney, Matthias Landgraf

**Affiliations:** University of Cambridge, Department of Zoology, Downing Street, Cambridge, CB2 3EJ, United Kingdom; HHMI Janelia Research Campus, Ashburn, VA, 20147, USA; Department of Biology, University of York, Heslington York YO10 5DD, United Kingdom

## Abstract

Neurons are inherently plastic, adjusting their structure, connectivity and excitability in response to changes in activity. How neurons sense changes in their activity level and then transduce these to structural changes remains to be fully elucidated. Working with the *Drosophila* larval locomotor network, we show that neurons use reactive oxygen species (ROS), metabolic byproducts, to monitor their activity. ROS signals are both necessary and sufficient for activity-dependent structural adjustments of both pre- and postsynaptic terminals and for network output, as measured by larval crawling behavior. We find the highly conserved Parkinson’s disease-linked protein DJ-1ß acts as a redox sensor in neurons where it regulates pre- and postsynaptic structural plasticity, in part via modulation of the PTEN-PI3Kinase pathway. Neuronal ROS thus play an important physiological role as second messengers required for neuronal and network tuning, whose dysregulation in the ageing brain and under neurodegenerative conditions may contribute to synaptic dysfunction.

## Introduction

Plasticity is fundamental to neuronal development and function. Changes in activity can trigger a raft of different responses, including changes to synaptic size, strength and connectivity, as well as to neuronal excitability (Davis and Müller, 2015; Keck et al., 2017; Wefelmeyer et al., 2015). ‘Hebbian’ plasticity mechanisms, such as long-term potentiation or depression, lead to altered transmission strength. These are balanced by homeostatic mechanisms, which are compensatory in nature and work toward maintaining neuronal or network activity within a set range. For example, blockade of excitatory inputs in neurons can induce a range of changes that are compensatory, including enlargement of both pre- and post-synaptic specializations, increasing synapse, axonal bouton and dendritic spine number as well as reducing dendritic spine elimination (Burrone et al., 2002; Kirov and Harris, 1999; Murthy et al., 2001; Zuo et al., 2005). How neurons monitor their activity levels is not known. Models of action potential firing rate homeostasis suggest that neurons measure the frequency of action potential firing and use homeostatic plasticity mechanisms to maintain this within their set point range (Davis, 2006; Turrigiano, 2012). In support of this view, recent studies in the mammalian visual system demonstrated that following dramatic activity disturbances, such as monocular occlusion, different types of neurons do indeed return to their original cell type-specific firing rate (Hengen et al., 2013; 2016).

An important second messenger in the regulation of activity-dependent plasticity is calcium, whose intracellular concentration correlates with neuronal activity patterns. Calcium regulates cellular and synaptic changes through a raft of signaling pathways, cytoskeletal and transcriptional effector proteins, including calcium/calmodulin-dependent protein kinase II (CaMKII), the phosphatase calcineurin and cyclic nucleotide signaling pathways (Vonhoff and Keshishian, 2017).

Here, we considered whether neurons might also monitor their activity levels by utilizing other signals. Neuronal activity levels correlate with energetic demand, suggesting that signals associated with energy metabolism could in principle provide a proxy measure for activity (Attwell and Laughlin, 2001; Hallermann et al., 2012; Zhu et al., 2012). We therefore focused on reactive oxygen species (ROS), which are formed as obligate byproducts of mitochondrial respiratory ATP synthesis; ROS form by ‘leakage’ of the electron transport chain, which leads to the generation of superoxide anions (O_2_-), and hydrogen peroxide (H2O2) (Halliwell, 1992). Intensively studied as destructive agents when at high concentration in the context of ageing and neurodegenerative conditions, in healthy cells ROS are necessary for many physiological processes, including growth factor signaling (Finkel, 2011). In cultured neurons, mitochondrial O_2_-production can be triggered by field stimulation (Hongpaisan et al., 2004; 2003). Increase in neuronal ROS levels were also reported following NMDA receptor stimulation, reportedly generated by mitochondria (Bindokas et al., 1996; Dugan et al., 1995) and NADPH oxidases (Brennan et al., 2009). In hippocampal and spinal cord slices, ROS were found sufficient and necessary for inducing ‘Hebbian’ forms of plasticity (LTP) (Kamsler and Segal, 2003a; 2003b; Klann, 1998; Knapp and Klann, 2002; Lee et al., 2010); and mice that over-express superoxide dismutase showed defects in hippocampal LTP and learning paradigms (Gahtan et al., 1998; Levin et al., 1998; Thiels et al., 2000).

To explore whether neurons might use metabolic ROS as signals for activity-induced structural plasticity, we used as a model the motor system of the fruitfly larva, *Drosophila melanogaster*, which is composed of individually identifiable motoneurons located in the ventral nerve cord and their specific target muscles in the periphery (Kohsaka et al., 2012). We had previously shown that ROS can regulate neuromuscular junction (NMJ) structure (Milton et al., 2011). Here, we show that the locomotor network of the *Drosophila* larva undergoes homeostatic adjustment in response to prolonged overactivation, as measured by larval crawling, its physiological output. This homeostatic adjustment of the network includes and requires structural plasticity of synaptic terminals. We show that ROS signaling, H2O2 in particular, is necessary and sufficient for this activity-induced synaptic terminal plasticity, of both presynaptic NMJ terminals and postsynaptic dendritic arbors within the central nervous system (CNS). As a sensor for activity-generated ROS we identified the conserved redox sensitive protein DJ-1ß, a homologue of vertebrate DJ-1 (PARK7) (Meulener et al., 2005). DJ-1ß genetically interacts with the phosphatase and tensin homolog (PTEN) and upon ROS mediated oxidation disinhibits PI3kinase signaling at the presynaptic NMJ, thus promoting synaptic terminal growth, as seen with activity-induced potentiation. Strong over-activation on the other hand leads to DJ-1ß-dependent compensatory adjustments of reduced release sites presynaptically and smaller dendritic input arbors.

## Results

### Induction of prolonged activity leads to homeostatic adjustment in the larval

#### *Drosophila* motor network

Neural circuits are inherently plastic and capable of adjusting their output with behavioral need. Our aim was to investigate mechanisms that allow neurons to sense changes in their activity levels and in response mediate adaptive structural adjustments. As an experimental model we used the locomotor network of the *Drosophila* larva, whose motoneurons grow dramatically over the course of larval life and are known to be plastic (Budnik et al., 1990; Davis, 2006; Schuster et al., 1996; Tripodi et al., 2008; Zito et al., 1999; Zwart et al., 2013). A simple way of manipulating activity in the locomotor network of *Drosophila* larvae is by changing ambient temperature (Dillon et al., 2009; Sigrist et al., 2003; Zhong and Wu, 2004). As previously documented (Sigrist et al., 2003; Zhong and Wu, 2004), we found that network activity increased upon *acutely* shifting animals to higher ambient temperatures (e.g. from 25°C to 29°C or 32°C), resulting in increased larval crawling speeds (Figure 1A). In contrast, following *prolonged* exposure to higher temperatures (rearing at 29°C and 32°C) we found that larvae crawled at the same speed as controls kept at 25°C (grey horizontal dotted line in Figure 1A), suggestive of homeostatic network adjustment. Mechanistically, neurons might compensate for the increased network drive that is the result of higher temperatures by reducing their individual excitability and/or synaptic input, thus returning motor output to the default crawling speed. In agreement with this model, when 29°C and 32°C-conditioned larvae were *acutely* shifted to a lower temperature of 25°C they displayed reduced crawling speed (green data in Figure 1a), with 32°C-adjusted larvae crawling significantly slower than 29°C -adjusted animals.

**Figure 1:**
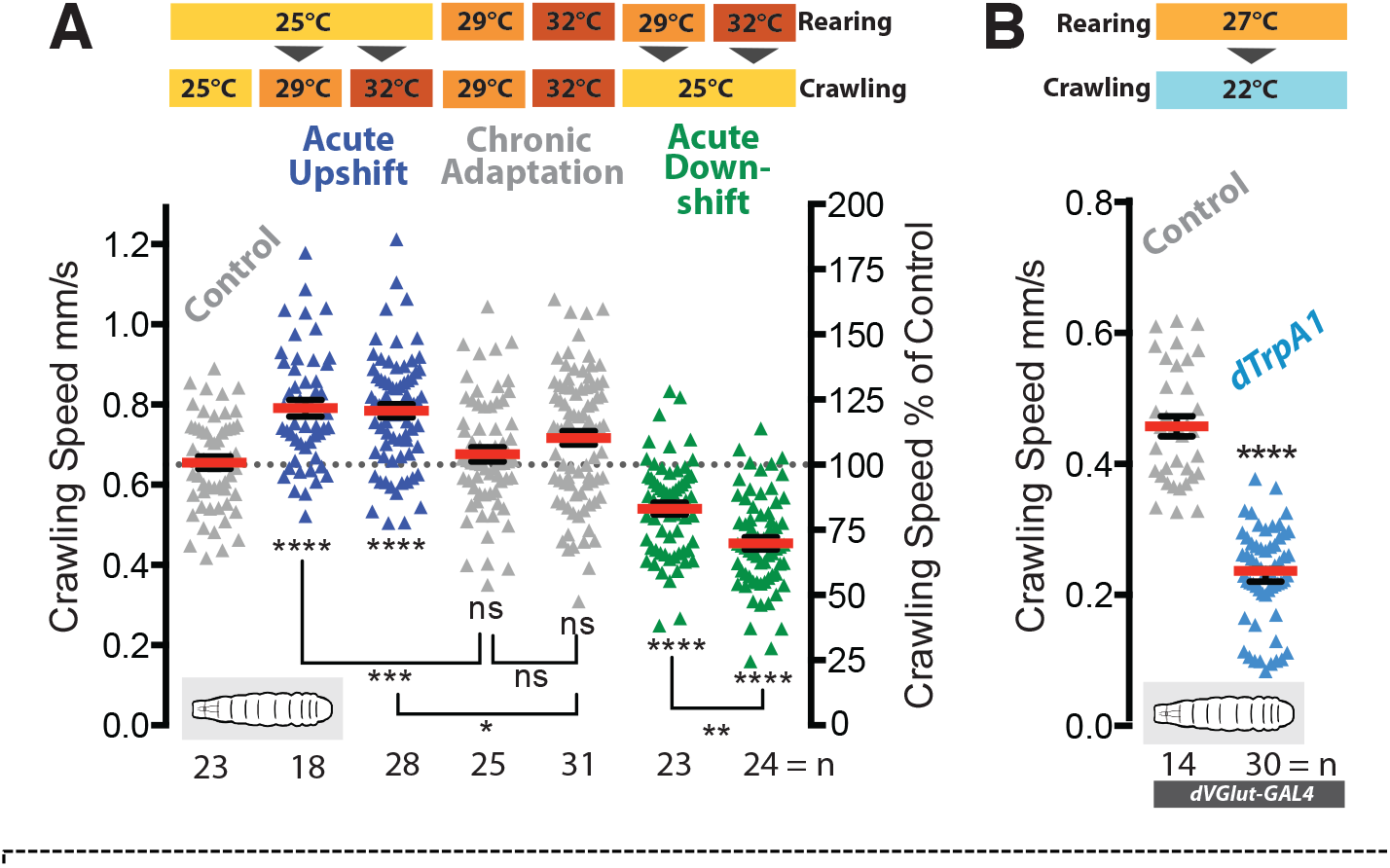
Adaptive behavioral plasticity is response to chromic locomotor over-activation. (**A**) Larval motor network activity, assayed by crawling speed 72hrs after larval hatching (ALH), increases in response to acute temperature upshifts (blue) in wild type larvae (Oregon R = control in all conditions). In contrast, with prolonged exposure (grey) to elevated temperatures (29°C or 32°C) the motor network adapts homeostatically generating the same crawling speed as 25°C reared controls. This adaptation is further revealed by acute temperature downshifts (green). Each data point represents crawling speed from an individual uninterrupted continuous forward crawl, n = specimen replicate number, up to 3 crawls assayed for each larva. Mean +/-SEM, ANOVA, ns = not significant, *P<0.05, **P<0.01, ***P<0.001, ****P<0.0001 (B) Prolonged over-activation targeted to motoneurons (*dVGlut-GAL4*; *UAS-dTrpA1*) also leads to adaptation with reduced crawling speed (dTRPA1 channels open at 27°C, closed at 22°C). Mean +/-SEM, control is *dVGlut-GAL4 / +.* ****P<0.0001 students t test, n = replicate number.

To test this further, we selectively over-activated the glutamatergic motoneurons by mis-expression of the warmth-gated cation channel dTrpA1 and rearing these animals at 27°C, a temperature that robustly activates dTrpA1-expressing neurons (Hamada et al., 2008; Pulver et al., 2009). Upon *acute* removal of this motoneuron over-stimulation (by shifting to 22°C, where the dTrpA1 channel is closed) larval crawling speed reduced significantly relative to non-dTrpA1 expressing controls (Figure 1B). Thus, prolonged increase of network drive leads to homeostatic adjustment that returns network output, and thus larval crawling speed, to a default set point. Cell type-selective manipulations of the motoneurons, which constitute the output of the motor network, show that they are major contributors to this adjustment.

### Structural plasticity of synaptic terminals is regulated by activity

We then asked what structural changes within the network might be associated and potentially responsible for these homeostatic behavioral adjustments. To this end, we focused on two well characterized motoneurons, ‘aCC’ and ‘RP2’ (Choi et al., 2004; Landgraf et al., 2003). Their presynaptic neuromuscular junction (NMJ) terminals consist of varicose compartments (boutons) strung across the muscle surface, each containing multiple synapses in the form of active zones (Van Vactor and Sigrist, 2017) (Figure 2A). We found that increases in motor network activity stimulated by higher ambient temperatures (raising animals at 29°C or 32°C) led to more, albeit smaller boutons at the NMJ by the wandering third instar larval stage, 100 hrs after larval hatching (ALH) (Figure 2C, grey data) (Sigrist et al., 2003; Zhong and Wu, 2004). To exclude non-specific effects, we targeted over-activation selectively to the aCC and RP2 motoneurons, via expression of the warmth-gated ion channel dTrpA1, and found similar presynaptic structural adjustments (Figure 2B & C, blue data; see also Figure 1 – supplement 1). While bouton number positively correlates with increases in motoneuron activity, quantification of synapse number, measured by quantification of active zones, describes a more complex relationship. As previously published, a moderate increase in activity (rearing larvae at 29^o^C) causes more boutons and also more active zones to be formed, thus potentiating transmission at the NMJ (Sigrist et al., 2003). In contrast, further increases in network activity, as effected by rearing larvae at 32°C, or cell-specific dTrpA1-mediated motoneuron activation led to progressive active zone reductions, consistent with a homeostatic response (Figure 2D, E).

**Figure 2:**
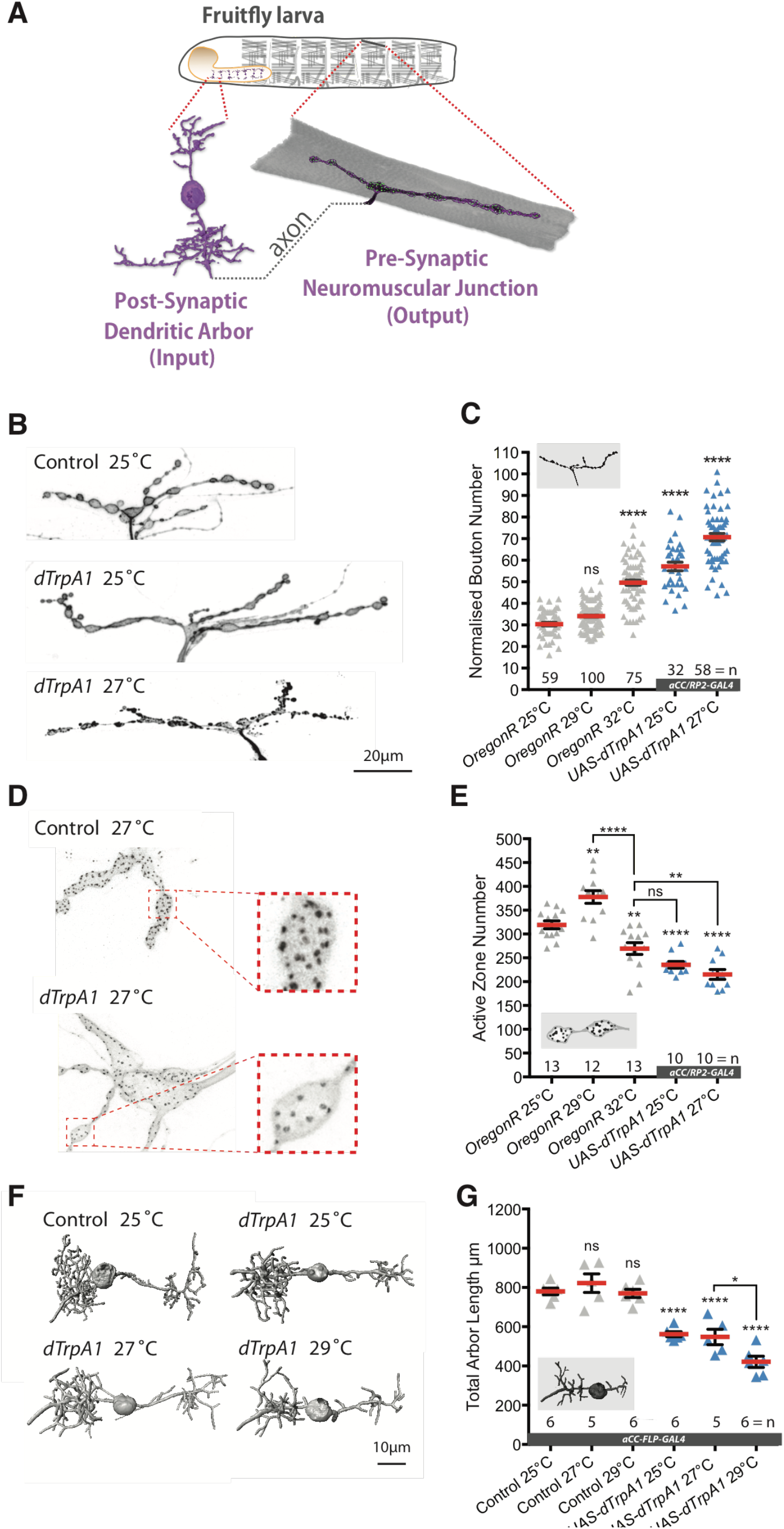
Adaptive structural synaptic plasticity at motorneuron input and output terminals in response to increased neuronal activity. (**A**) Graphical illustration of a stereotypical larval motorneuron (MN), adapted from (Kohsaka et al., 2012). Pre-motor interneurons make synaptic connections with the MN dendritic arbor (input) in the larval ventral nerve cord (equivalent of mammalian spinal cord). The MN extends an axonal projection into the periphery where it connects with a target muscle via an NMJ, characterized by varicose swellings (boutons) each containing multiple individual neurotransmitter release sites (active zones). (**B** and **C**) Representative images of muscle DA1[muscle 1] NMJs from 3^rd^ instar larvae (100hrs ALH). Dot-plot quantification shows NMJ bouton number increases in response to systemic and cell-specific activity increases. (**D** and **E**) Active zone number increases following low-level over-activation (29°C), but progressively reduces upon stronger over-activation. (**F** and **G**) Digital reconstructions and dot plots show that over-activation leads to reduced total dendritic arbor length of aCC motoneurons (24hrs ALH). ‘*aCC/RP2-GAL4’* expresses GAL4 in all, ‘*aCC-FLP-GAL4’* in single aCC and RP2 motoneurons (see Online Methods for details); ‘Control’ in (F + G) is *aCC-FLP-GAL4* alone. Mean +/-SEM, ANOVA, ns = not significant, *P<0.05, **P<0.01, ***P<0.001, ****P<0.0001, n = replicate number. Comparisons with control are directly above data points.

We then turned to look at the input arbor of the motoneuron, branched dendrites on which the pre-motor interneurons form connections (Baines et al., 1999; Schneider-Mizell et al., 2016; Zwart et al., 2013). Using a stochastic labeling strategy within a small subset of motoneurons (aCC-FLP-GAL4) (Ou et al., 2008), we selectively targeted GAL4 and dTrpA1 expression to individual aCC motoneurons. Following confocal imaging and digital reconstructions of the arbors (Evers et al., 2005; Schmitt et al., 2004), morphometric analysis revealed that the size of the postsynaptic dendritic arbor decreased with rising levels of temperature-gated dTrpA1 activity (Figure 2F, G). Because in these motoneurons dendritic length correlates with input synapse number and synaptic drive (Tripodi et al., 2008; Zwart et al., 2013), we interpret this structural adjustment of the dendrites as homeostatic.

Thus, while a moderate increase in neuronal activity can lead to potentiation of the presynaptic motoneuron terminal by increasing bouton and synapse number (Ataman et al., 2008; Piccioli and Littleton, 2014; Sigrist et al., 2003), stronger over-activation leads to structural adjustments at both input and output terminals of the motorneuron that we interpret as compensatory; smaller dendritic arbors being associated with less synaptic drive (Zwart et al., 2013) and fewer presynaptic release sites at the NMJ reducing transmission to the muscle.

### Activity generated ROS regulate structural plasticity at synaptic terminals

Because neural activity is energetically demanding (Attwell and Laughlin, 2001; Hallermann et al., 2012; Zhu et al., 2012) we wondered whether neurons might monitor activity levels through metabolic signals. Others previously reported that neuronal over-activation *in vitro* can lead to a decrease in the ATP:ADP ratio (Tantama et al., 2013) and an increase in mitochondrial ROS (Hongpaisan et al., 2004). Moreover, at the *Drosophila* NMJ oxidative stress has been shown to regulate NMJ structure (Milton et al., 2011). To ask whether ROS signaling might be associated with activity-dependent synaptic terminal growth, we expressed the mitochondrion-targeted ratiometric ROS reporter *UAS-mito-roGFP2-Orp1* in aCC and RP2 motoneurons (Gutscher et al., 2009). This reports a titratable neuronal activity-dependent increase in mitochondrial ROS (Figure 3A).

**Figure 3:**
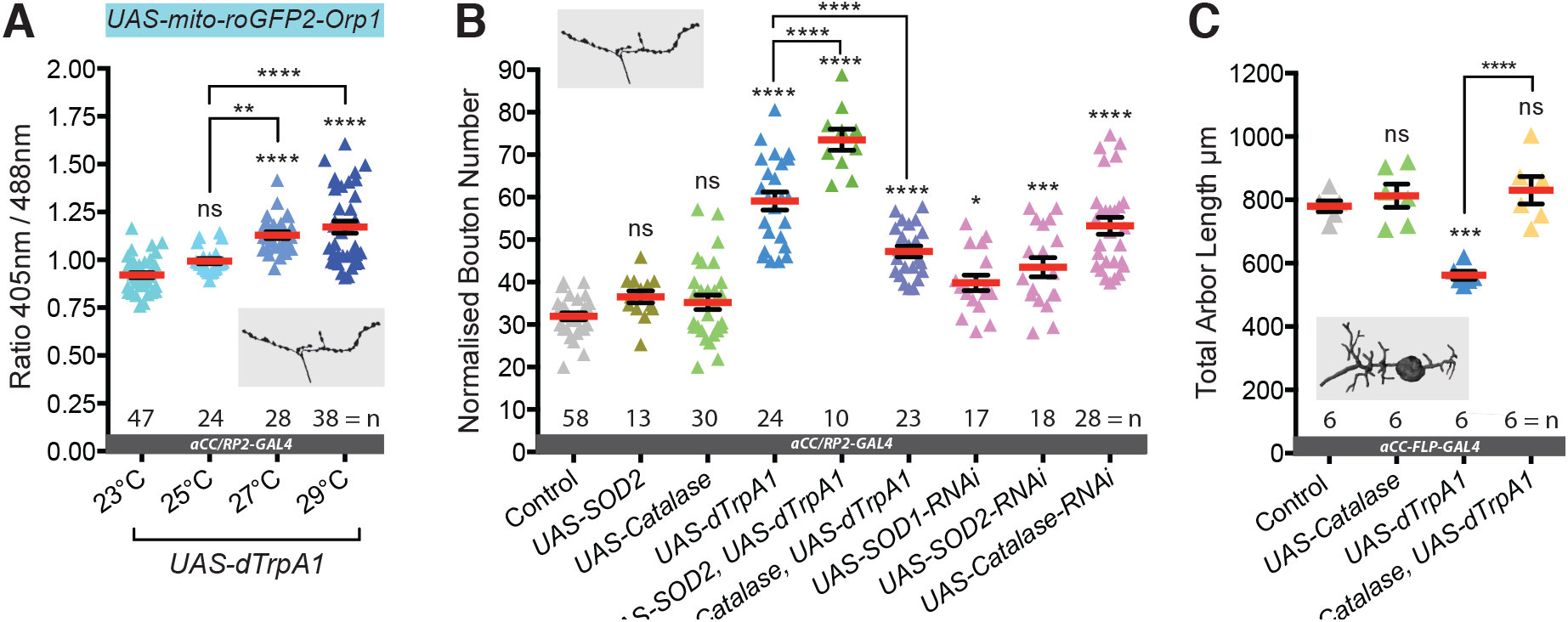
Neuronal activation leads to generation of synaptic ROS that regulate structural plasticity at input and output terminals. (**A**) Elevated neuronal activity increases mitochondrial ROS production at NMJs. Dot plots of mitochondrion-targeted ratiometric H_2_O_2_ sensor (*UAS-mito-roGFP2-Orp1* (Gutscher et al., 2009)) in wandering 3^rd^ instar larval NMJs (100hrs ALH) at 23˚C (control, dTrpA1 inactive), 25˚C (moderate activity, ~10Hz), 27˚C (strong activity, ~20Hz) and 29˚C (very strong excitation, ~40Hz). (**B**) Bouton number at the NMJ is increased by *UAS-dTrpA1*-mediated over-activation. This is exacerbated by co-expression of *UAS-SOD2* (converts O_2_- to H_2_O_2_) and rescued by the H_2_O_2_ scavenger *UAS-Catalase*. Cell-specific ROS elevation by scavenger knockdown is sufficient to induce NMJ elaboration. *aCC/RP2-GAL4*, ‘Control’ is *aCC/RP2-GAL4* alone. (**C**) Total dendritic arbor length is reduced by single cell over-activation, but rescued by co-expression of the H_2_O_2_ scavenger *UAS-Catalase* (aCC motoneurons, 24hr ALH). *aCC-FLP-GAL4*, ‘Control’ is *aCC-FLP-GAL4* alone. Larvae reared at 25°C. Mean +/-SEM, ANOVA, ns = not significant, *P<0.05, **P<0.01, ***P<0.001, ****P<0.0001, n = replicate number.

We therefore hypothesized that mitochondrially derived ROS might provide a readout of neuronal activity and regulate synaptic structural plasticity. To test this hypothesis, we selectively increased neuronal activity in the aCC and RP_2_ motoneurons while at the same time over-expressing the ROS scavenging enzymes Superoxide Dismutase 2 (SOD_2_, which catalyses O_2_- to H_2_O_2_ reduction) or Catalase (H_2_O_2_ into H_2_O and O_2_). Catalase co-expression significantly counteracted dTrpA1 activity-induced bouton addition at the NMJ (Figure 3B) and rescued motoneuron dendritic arbor size in the CNS (Figure 3C). SOD_2_ co-expression on the other hand enhanced dTrpA1-mediated NMJ elaboration, presumably by potentiating conversion of neuronal activity-induced O_2_- into H_2_O_2_ (Figure 3B). We then confirmed that ROS are also sufficient to invoke structural plasticity in the absence of neuronal activity manipulation through cell-specific RNAi knock down of SOD1, SOD2 and Catalase (Figure 3B). All of these manipulations led to NMJ growth phenotypes with increased bouton number that are similar to those produced by neuronal over activation. Based on these results, we implicate ROS, specifically H2O2, in the observed activity-dependent structural plasticity.

### DJ-1ß acts as a ROS sensor in neurons

We then asked how neurons might sense changes in activity-induced ROS levels. ROS are known to post-transcriptionally modify many different proteins, principally on cysteine residues, including transcription factors, cell adhesion molecules and phosphatases (for review see (Milton and Sweeney, 2012)). Following a literature review, we focused on DJ-1ß, the fly ortholog of DJ-1 (PARK7), as a candidate ROS sensor. *DJ-1* codes for a highly conserved, ubiquitously expressed redox-sensitive protein, that protects against oxidative stress and regulates mitochondrial function (Ariga et al., 2013; Nagakubo et al., 1997). A mutant allele is also linked to a rare form of familial Parkinsonism (Bonifati et al., 2003). *DJ-1* null mutant adult flies had been reported to be more sensitive to oxidative stress, induced by exposure to paraquat and H_2_O_2_ (Meulener et al., 2005). We found that *DJ-1ß* null mutant (*DJ-1ß*^*Δ93*^) larvae develop normally and have structurally normal NMJs (Figure 4A and Figure 4-supplement 1). What is more, systemic loss of DJ-1ß (*DJ-1ß*^*Δ93*^ null mutants) completely rescues NMJ bouton addition phenotypes normally caused by pharmacologically induced oxidative stress (Milton et al., 2011) (Figure 4A). Loss of DJ-1ß also significantly rescues NMJ bouton phenotypes induced by cell type-specific expression of the ROS generator Duox or dTrpA1-mediated over-activation (Figure 4A). *DJ-1ß*^*Δ93*^ mutant larvae failed to produce the presynaptic bouton and active zone addition characteristic of NMJ potentiation as seen in controls following periods of temperature-stimulated (29^o^C) increases in locomotor activity (Sigrist et al., 2003) (Figure 4 – supplement 2). Motoneuron-targeted expression of a dominant-acting mutant form of DJ-1ß that is non-oxidizable at the conserved cysteine 104 (DJ-1ß^C104A^) abrogated dTrpA1 activity-mediated NMJ structural adjustment, both with respect to bouton number (Figure 4A) and active zone number (Figure 4B). Looking at structural plasticity of the postsynaptic dendritic arbor, we found that halving the *DJ-1ß* copy number (in *DJ-1ß*^*Δ93*^*/+* heterozygotes) was sufficient to significantly suppress dTrpA1-mediated reductions in dendritic arbor size (Figure 4C).

**Figure 4:**
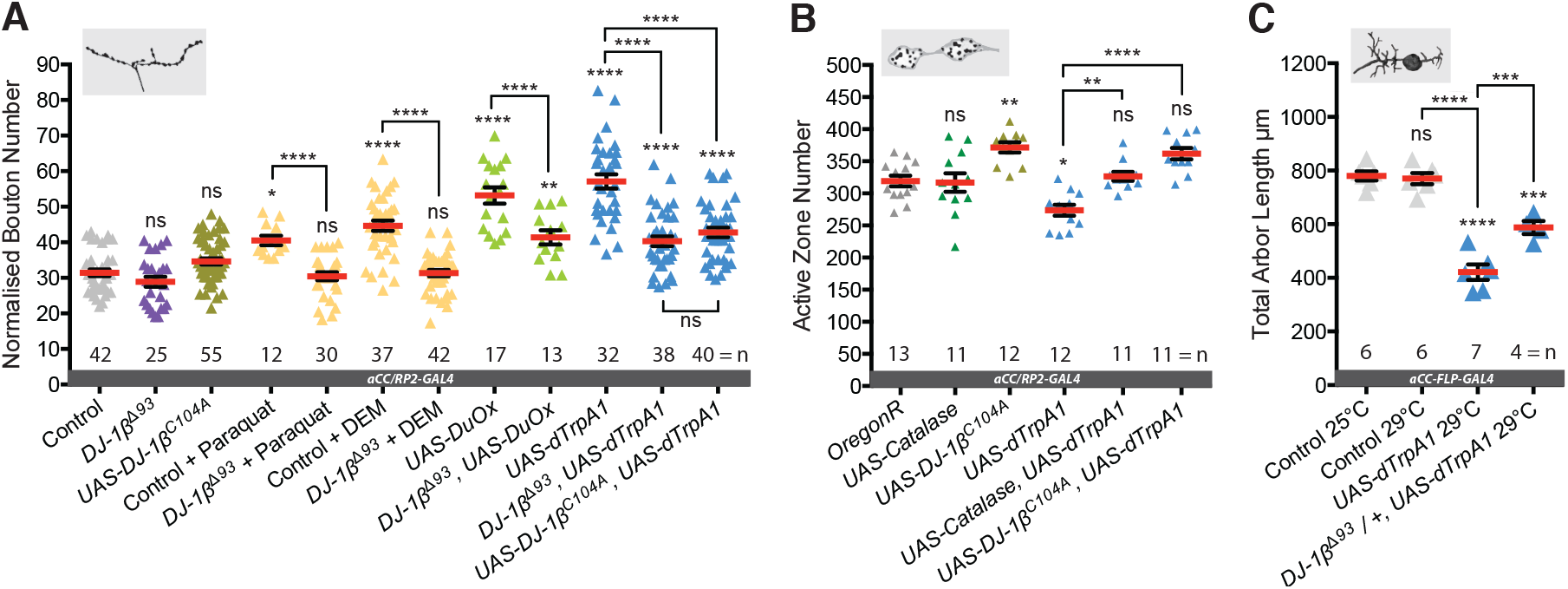
*DJ-1β* senses ROS and regulates activity-induced neural plasticity. (**A**) *DJ-1β* is required for ROS and neuronal activity-induced NMJ elaboration (100hrs ALH). Larvae reared at 25°C. (**B**) Cell-specific expression of *DJ-1β*^*C104A*^, non-oxidizable on conserved cysteine C104, prevents activity-induced reduction of active zone number. *aCC/RP2-GAL4*, ‘Control’ is *aCC/RP2-GAL4* alone. Larvae reared at 25°C. (**C**) Activity generated ROS sensing is dose sensitive. Removal of one copy of *DJ-1β* (in *DJ-1β*^*Δ93*^ */ +* heterozygotes) is sufficient to significantly rescue activity-induced reduction of total dendritic arbor length of motoneurons in 24hr ALH larvae. *aCC-FLP-GAL4*, ‘Control’ is *aCC-FLP-GAL4* alone.

In summary, these data show that DJ-1ß is necessary for sensing activity-induced ROS and for implementing structural changes at both pre- and postsynaptic terminals in response. At the presynaptic NMJ, DJ-1ß appears to be required in motoneurons for at least two kinds of activity regulated structural plasticity: following mild over-activation, the addition of both boutons and active zones, which are associated with potentiation (Sigrist et al., 2003); and following stronger over-activation, reduction of active zone number, which we interpret as an adaptive adjustment.

### Structural plasticity of synaptic terminals is required for homeostatic adjustment of locomotor behavior

We then asked whether this ROS - DJ-1ß mediated structural plasticity of synaptic terminal growth was required for the homeostatic adjustments of crawling speed, which ensues after prolonged over-activation. To answer this question, we targeted expression of the dominant acting non-oxidizable DJ-1ß^C104A^ to motoneurons (and other glutamatergic cells, using DvGlut-T2A-GAL4). We then tested the behavior of these animals for adjustment in response to *chronic* temperature-induced elevation of motor network activity. Our previous experiments showed that expression of non-oxidizable DJ-1ß^C104A^ in motoneurons does not lead to abnormal bouton number or the decrease in active zone number normally caused by over-activation (see Figures 4A & 4B). Expression of non-oxidizable DJ-1ß^C104A^ in motoneurons per se did not alter larval crawling speed at the control temperature of 25^o^C. However, when rearing these larvae at 32^o^C, which is associated with elevated motor network activation, unlike controls they failed to homeostatically adjust toward the default crawling speed set point (Figure 5). Consequently, such larvae reared at elevated temperatures (29^o^C or 32^o^C) also responded less strongly than controls to acute temperature downshifts (Figure 5). These data suggest that activity-induced neuronal plasticity, implemented through the ROS sensor DJ-1ß, is necessary for activity-directed homeostatic adjustment of larval locomotor behavior.

**Figure 5:**
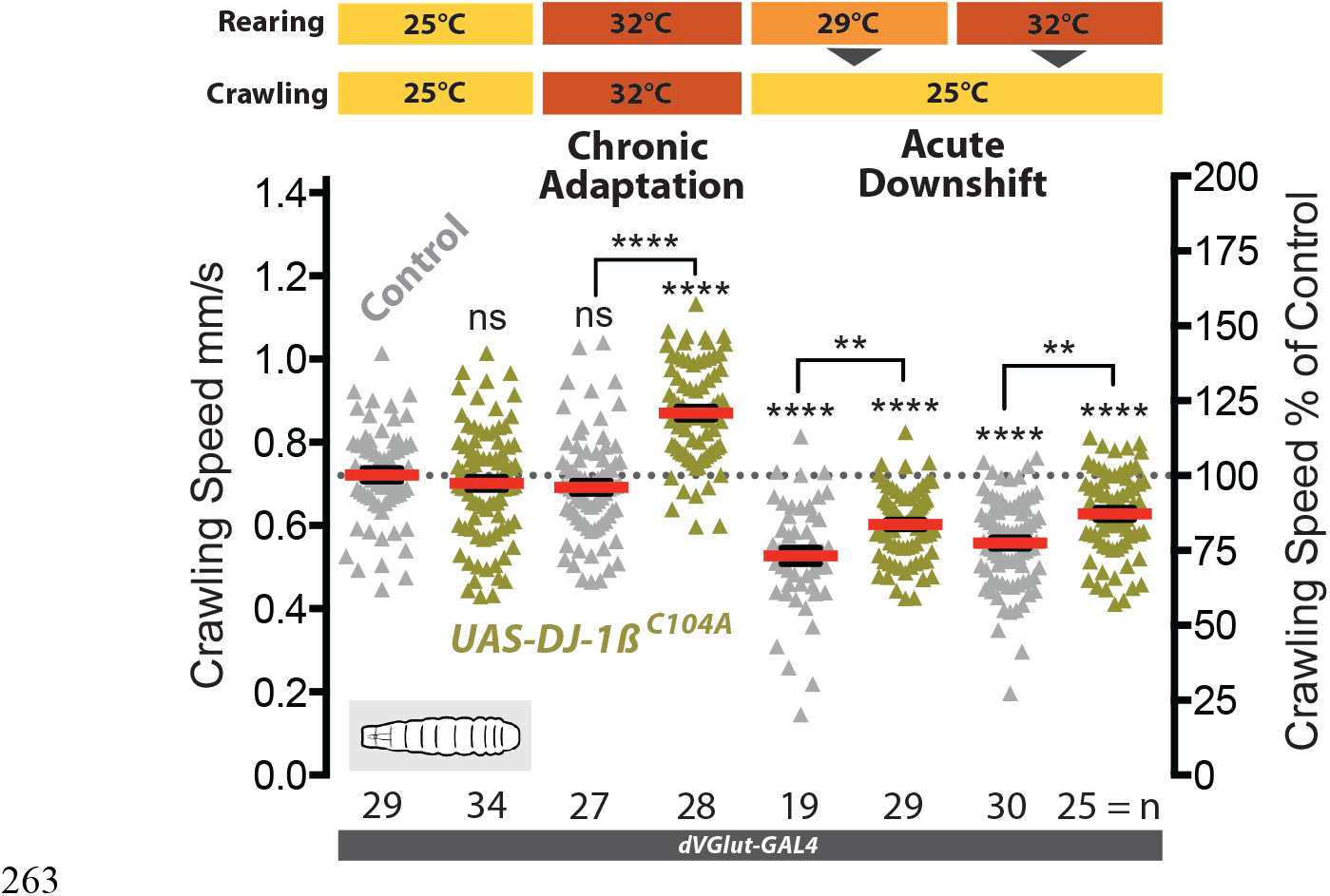
*DJ-1β* is required in motoneurons for behavioral plasticity in response to chronic locomotor over-activation. Larvae with expression of *UAS-DJ-1β*^*C104A*^ targeted to motoneurons (*dVGlut-GAL4*) are unable to adapt motor network output (crawling speed) to elevated rearing temperatures. Control is *dVGlut-GAL4* alone. Each data point represents crawling speed from an individual uninterrupted continuous forward crawl, n = specimen replicate number, up to 3 crawls assayed for each larva. Mean +/-SEM, ANOVA, ns = not significant, **P<0.01, ****P<0.0001, n = replicate number.

### PTEN and PI3K are downstream effectors of the DJ-1ß ROS sensor

Next, we looked for downstream effector pathways responsible for implementing activity and ROS-dependent structural plasticity. DJ-1 is a known redox-regulated inhibitor of Phosphatase and Tensin homologue (PTEN) and thus a positive regulator of PI3Kinase signaling (R. H. Kim et al., 2005; Y.-C. Kim et al., 2009); and PI3Kinase is a known positive regulator of NMJ development (Jordán-Álvarez et al., 2012; Martín-Peña et al., 2006). To test whether DJ-1ß – PTEN interactions mediate ROS-dependent NMJ adjustments, we performed genetic interaction experiments in the context of a DEM (oxidative stress) dose-response curve (Figure 6A). Focusing on the NMJ, we found that in controls bouton number increases linearly with exposure to increased DEM concentrations, peaking at 15mM DEM (Figure 6A). Removing one copy of *DJ-1ß* (*DJ-1ß*^*Δ93*^*/+*) was sufficient to suppress DEM-induced increases in bouton number. In contrast, *PTEN*^*CO76*^*/+* heterozygous larvae showed increased sensitivity to DEM, with a left-shifted dose response curve. Larvae heterozygous for both *DJ-1ß* and *PTEN* (*PTEN*^*CO76*^*/+; DJ-1ß*^*Δ93*^*/+*) were more sensitive to DEM at higher concentrations than *DJ-1ß*^*Δ93*^*/+* heterozygotes. These genetic interactions agree with previous studies (R. H. Kim et al., 2005) and complement biochemical data that showed H_2_O_2_-oxidized DJ-1ß binds to and inhibits PTEN (Y.-C. Kim et al., 2009).

**Figure 6:**
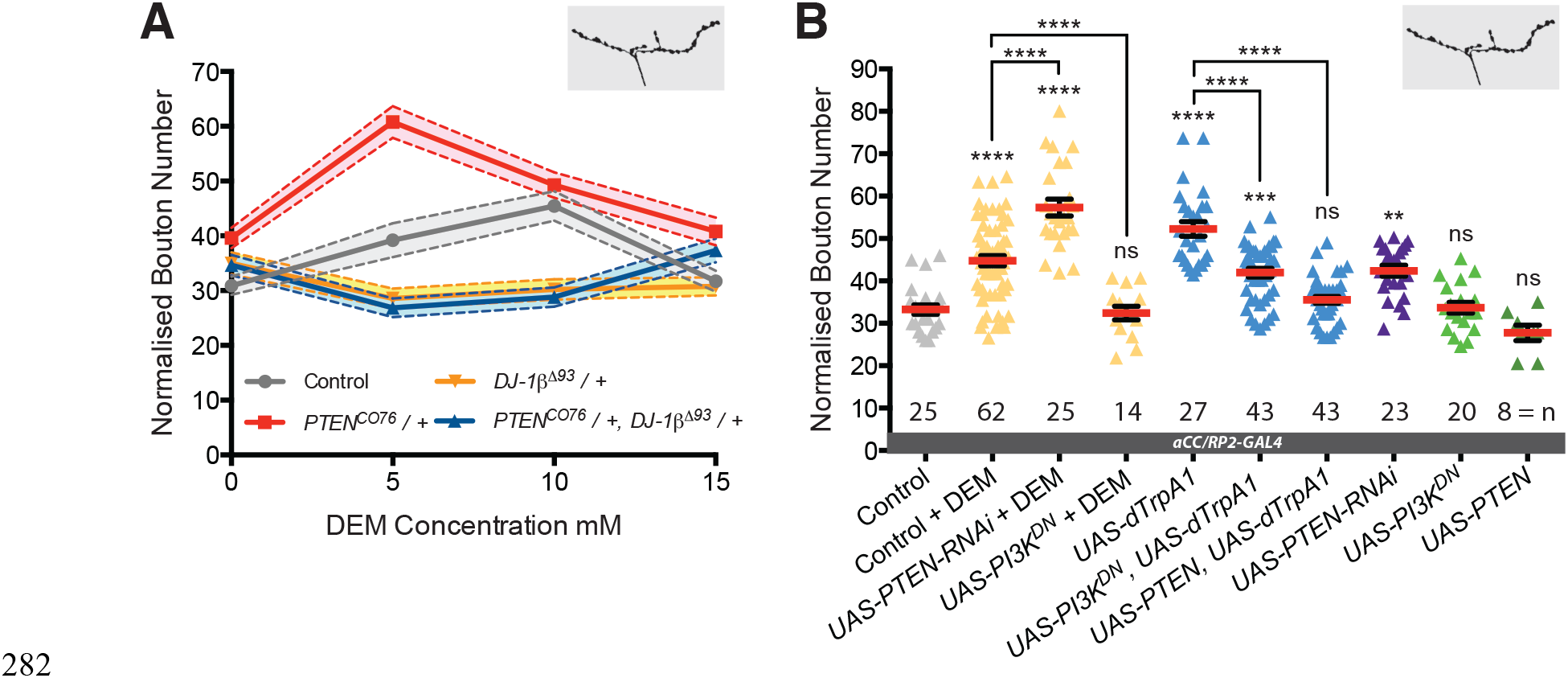
*DJ-1β* signals via PTEN and PI3Kinase to regulate ROS and activity-induced NMJ elaboration. (**A**) *DJ-1β* and PTEN genetically interact to regulate systemic ROS-induced NMJ elaboration. NMJ bouton number varies with ROS (DEM) levels (grey data). Removal of one copy of *PTEN* sensitizes (red) while heterozygosity for *DJ-1ß* desensitizes NMJs to ROS levels (yellow), partially restored in double heterozygotes (blue). Dashed boundaries indicate 95 % confidence intervals (n≥38). (**B**) Systemic ROS and activity-induced NMJ structural adjustments require PTEN and PI3Kinase signaling. Over-expression of the PI3Kinase antagonist PTEN or a dominant negative PI3Kinase form abrogates activity-induced NMJ elaboration. *aCC/RP2-GAL4*, ‘Control’ is *aCC/RP2-GAL4* alone. Mean +/-SEM, ANOVA, ns = not significant, **P<0.01, ***P<0.001, ****P<0.0001, n = replicate number.

To further test specificity, we manipulated PTEN and PI3Kinase activities in single cells. Targeted knock-down of PTEN in motoneurons sensitized these neurons to ROS, exacerbating the DEM-induced bouton addition phenotype (Figure 6B). In contrast, overexpression of PTEN or mis-expression of a dominant negative form of PI3Kinase significantly reduced NMJ elaboration normally caused by DEM exposure or dTrpA1-mediated neuronal activity increase (Figure 6B). Together these interactions suggest that PTEN and PI3Kinase act downstream of DJ-1ß and neural activity-generated ROS; and that oxidation of DJ-1ß, through PTEN inhibition, facilitates a rise in PI3Kinase / PIP3 signaling, which in turn mediates at least part of the structural synaptic terminal plasticity (Figure 7).

**Figure 7:**
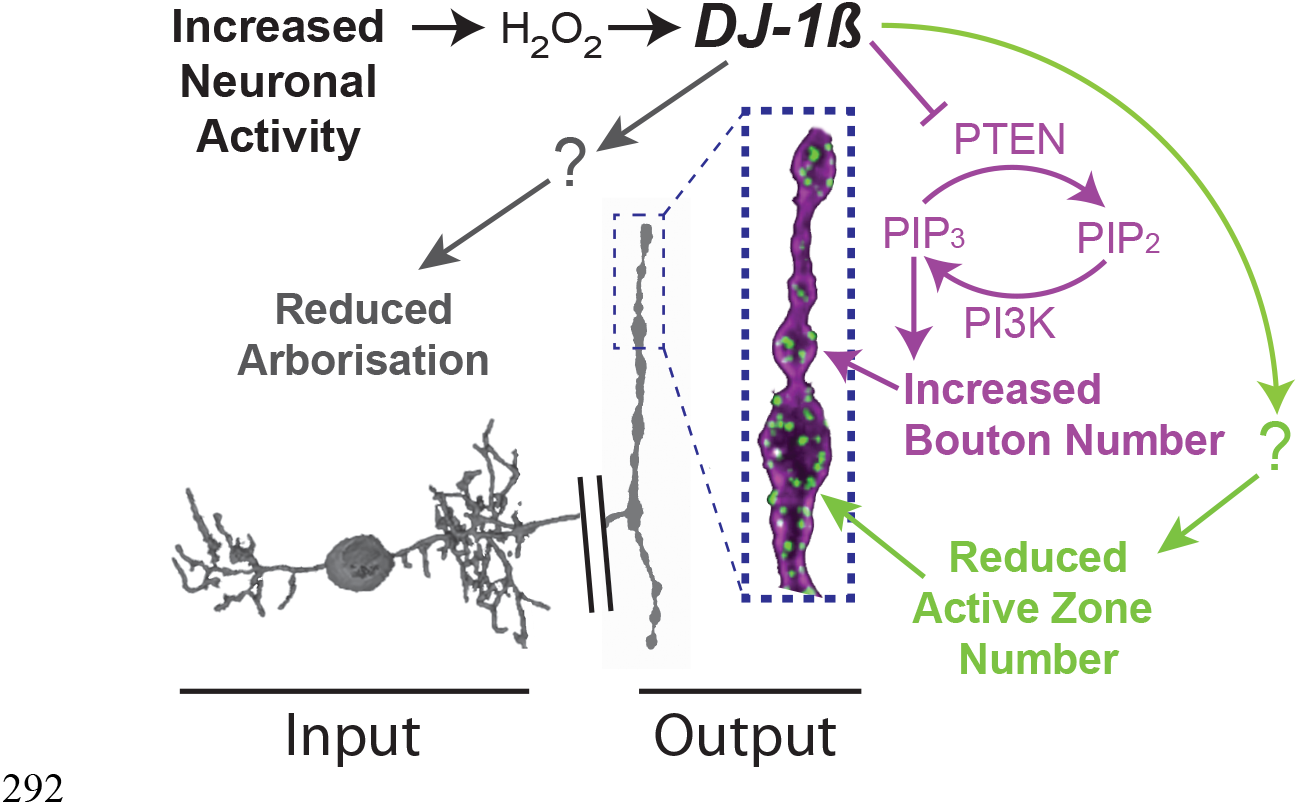
Model summary. *DJ-1β* is a redox signaling hub that coordinates structural synaptic plasticity at motoneuron synaptic input and output terminals. Activity-induced ROS oxidize *DJ-1β*, leading to PTEN inhibition and thus to a gain in PI3Kinase signaling, which regulates activity-induced NMJ elaboration of boutons and active zones. At higher activity/ROS thresholds additional, yet to be defined, pathways downstream of *DJ-1β* are activated, implementing adaptive reductions of active zones at the NMJ and dendritic arbor length in the CNS.

## Discussion

Our aim was to study mechanisms of structural plasticity in relation to neuronal activity in a motor network. To do so we used as a model the *Drosophila* larva and focused on major effectors of motor circuit output: motoneurons, their dendrites in the CNS and presynaptic terminals at peripheral neuromuscular junctions. We have identified a novel pathway by which these neurons sense changes in activity through measuring increases in activity-generated ROS, which act as second messengers for structural adjustments at both dendritic input and presynaptic output terminals. Our data further suggest that this activity-regulated structural plasticity is instrumental for the homeostatic adjustment of larval locomotor network activity and thus crawling behavior.

### Homeostatic adjustment of the motor network allows animals to maintain a dynamic range of crawling speed

To investigate forms and mechanisms of structural plasticity that neurons undertake while adapting to altered activity levels, we took advantage of *Drosophila* motor network activity being readily manipulated through changes in ambient temperature. We found that while acute increases in ambient temperature from 25^o^C to 29^o^C or above quickly push larvae to their maximum crawling speed (approx. 0.8mm/sec), prolonged exposure leads to the network adjusting homeostatically toward a default crawling speed (approx. 0.65mm/sec) (Figure 1A). This adaptation might overall be energetically more favorable, and it allows larvae to stay within their dynamic range of behavioral responses to relative changes regardless of absolute ambient temperature.

Cell type-specific over-activation of the glutamatergic motoneurons (via dTrpA1) largely recapitulated the larval crawling phenotypes otherwise generated by systemic motor network overactivation. Conversely, blockade of activity-induced structural adjustment (through mis-expression of non-oxidisable DJ-1ß^C104A^) in the glutamatergic motoneurons significantly blocked homeostatic locomotor network adjustment to higher ambient temperatures (Figure 1B). The importance of the motoneurons in shaping much of the adaptive behavioral response is perhaps to be expected, since their dendrites are the final integrators on which all pre-motor inputs converge (Fushiki et al., 2016; Itakura et al., 2015; Kohsaka et al., 2014; Schneider-Mizell et al., 2016; Zwart et al., 2016). Systemic manipulations produce stronger behavioral phenotypes than those targeted to the glutamatergic neurons, suggesting, as would be expected, that other cells within the locomotor network are also subject to activity-regulated homeostatic plasticity (Figure 5). Interestingly, a recent study, which modeled different forms of plasticity in this motor network, concluded that during its development homeostatic mechanisms are better placed in achieving the formation of a stable balanced network producing robust activity patterns than Hebbian alternatives (Gjorgjieva et al., 2016).

### Structural adjustment coordination between pre- and postsynaptic terminals

Working with identified motoneurons we attempted to take an integrated view by relating structural plasticity across its pre-synaptic and dendritic terminals. At the presynaptic NMJ low-level increases of activity lead to potentiation, associated with increased numbers of active zones, elevated levels of the Brp/CAST/ELKS family active zone protein Bruchpilot (Brp) and the addition of boutons, varicose swellings containing multiple active zones (Cho et al., 2015; Sigrist et al., 2003; Vasin et al., 2014; Weyhersmüller et al., 2011).It is known that changes to the composition of the active zone cytomatrix (Davydova et al., 2014; Lazarevic et al., 2011; Matz et al., 2010) such as elevation of Brp levels can further aid transmission by increasing release probability (Peled et al., 2014; Weyhersmüller et al., 2011).

Strong over-activation, in contrast, though triggering further addition of boutons, leads to overall reductions in active zone number at the NMJ, which correlate with the severity of over-activation (Figure 2D, E). This parallels observations in the adult *Drosophila* visual system where increased activation also triggers a reduction of Brp levels in photoreceptor terminals (Sugie et al., 2015). We interpret this negative correlation between the degree of over-activation and the reduction in active zone number, and potentially Brp content, as a compensatory response. In agreement, blocking activity-induced structural adjustment in the motoneurons abrogates behavioral adaptation to prolonged over activation (Figure 5). Whether increased numbers of smaller boutons forms part of an adaptive response to over-activation or is anticipatory of future synapse strengthening remains to be established.

The input terminal of the neuron, the dendritic arbor in the CNS, responds to overactivation with corresponding reductions in dendritic length, which we previously found to correlate with reducing both input synapses and synaptic drive. We therefore interpret this activity driven structural adjustment of the dendrites also as compensatory (Tripodi et al., 2008; Zwart et al., 2013). In summary, we found that above threshold over-activation regimes lead to structural adaptive adjustments at both the postsynaptic input and presynaptic output terminals. We do not yet know whether these are regulated locally or globally coordinated by transcriptional changes; though at least for the presynaptic NMJ activity and ROS mediated structural changes have been reported as dependent on immediate early genes (AP-1) (Milton et al., 2011; Sanyal et al., 2002). Given the available evidence we would expect these structural adjustments to be linked to, and potentially preceded by compensatory changes in neuronal excitability (Baines et al., 2001; Davis, 2006; Davis et al., 1996; 1998; Driscoll et al., 2013; Frank et al., 2006; 2009; Gaviño et al., 2015; Giachello and Baines, 2016; Lin et al., 2012; Mee et al., 2004; Müller and Davis, 2012; O’Leary et al., 2013; Prinz, 2006; Prinz et al., 2004; Wang et al., 2014; Younger et al., 2013).

### ROS and redox sensitive DJ-1ß provide a readout of neuronal activity

Neurons have cell type-specific set points of average firing ranges to which they return following activity perturbations using a variety of homeostatic plasticity mechanisms (Hengen et al., 2016; 2013; Keck et al., 2017; 2013). The mechanisms that determine the homeostatic set point and allow neurons to sense when their activity deviates from such a set point are not yet known. Intracellular concentrations of calcium correlate with neuronal activity (Hardingham et al., 2001), making this versatile second messenger a prime candidate for providing a readout of activity and for regulating homeostatic plasticity (O’Leary et al., 2014). In this study we asked whether metabolically associated signals, such as ROS, might also be involved, as these could provide a proxy measure of neuronal activity (Attwell and Laughlin, 2001; Zhu et al., 2012). Over-activation measurably impacts on the ADP:ATP ratio (Tantama et al., 2013) and generates mitochondrial ROS. These can modulate calcium dependent signaling pathways, such as CamKII (Hongpaisan et al., 2004), and potentially also other ROS sensitive pathway components including IP3 and Ryanodine receptor channels or the protein phosphatase Calcineurin (Hidalgo and Arias-Cavieres, 2016). We found that a mitochondrially targeted ROS sensor (Albrecht et al., 2011; Gutscher et al., 2009) became progressively oxidized with increasing levels of neuronal over-activation (Figure 3A). Moreover, ROS were necessary for (and their elevation was also sufficient to mimic) implementing activity-induced structural plasticity at synaptic terminals (Figure 3B, C). Thus, our data show that ROS, particularly H_2_O_2_, are both necessary and sufficient for activity regulated structural plasticity. The source of the activity-generated ROS appears to be mitochondrial, potentially generated as a byproduct of increased ATP metabolism or triggered by mitochondrial calcium influx (Peng and Jou, 2010). However, other ROS generating systems could also be involved, notably NADPH oxidases, some of which are regulated by calcium (Kishida et al., 2006; 2005; Serrano et al., 2003; Tejada-Simon et al., 2005).

We discovered that in neurons the highly conserved protein DJ-1ß is critical for responding to activity-generated ROS with structural changes to synaptic terminals (Figure 4A, B, C), and propose that in neurons DJ-1ß acts as a redox sensor for activity-generated ROS. In agreement with this idea, DJ-1ß has been shown to be oxidized at the conserved cysteine residue C106 (C104 in *Drosophila*) by H_2_O_2_. Oxidation of DJ-1 leads to changes in DJ-1 function, including translocation from the cytoplasm to the mitochondrial matrix, which aids with protection against oxidative damage (Blackinton et al., 2009; Canet-Avilés and Wilson, 2004; Waak et al., 2009) and maintenance of ATP levels (Calì et al., 2015). We found that motoneurons are potently sensitive to DJ-1ß dosage with regard to their response to altered activity and ROS levels, and expression of mutant DJ-1ß^C104A^ in motoneurons blocks their structural plasticity response to DEM and over-activation. These observations suggest that DJ-1ß forms a critical part of the ROS sensing mechanism in neurons. It is therefore conceivable that different sensitivities to neuronal activity could in part be determined by cell type-specific levels of DJ-1ß and associated reducing mechanisms.

### DJ-1ß downstream pathways implement activity-regulated plasticity

Our data suggest that DJ-1ß forms part of a signaling hub required for at least two opposing types of structural plasticity that seem to be implemented by distinct pathways: potentiation in response to low-level increases in neuronal activation versus compensatory changes following stronger over-activation. In this study we demonstrated that DJ-1ß mediates the former by activation of PI3Kinase signaling following redox-regulated binding and inhibition of PTEN (Figure 4A) (R. H. Kim et al., 2005; Y.-C. Kim et al., 2009). PTEN and PI3Kinase are both intermediates of metabolic signaling pathways and regulators of synaptic terminal growth (Jordán-Álvarez et al., 2012; Martín-Peña et al., 2006). While activity and ROS levels correlate positively with PTEN/PI3Kinase mediated bouton addition at the NMJ, active zone number regulation is more nuanced; at a higher activity threshold other, yet to be identified effector pathways appear to be activated that reduce presynaptic active zone number (Figure 2D, E). Because this too is dependent on DJ-1ß we would expect this pathway also to be regulated by activity-generated ROS. Equally unresolved is the question of how postsynaptic dendritic arbor size in the CNS is regulated. PTEN/PI3Kinase signaling is an unlikely candidate as this normally stimulates neuronal and dendritic growth (Brierley et al., 2009).

### ROS as gatekeepers of activity-dependent synaptic structural plasticity

Synaptic plasticity is regulated by a number of signaling pathways, including Wnts (Budnik and Salinas, 2011), BMPs (Bayat et al., 2011; Berke et al., 2013), as well as PKA, CREB and AP-1 (Cho et al., 2015; Davis, 2006; Davis et al., 1998; 1996; Davis and Müller, 2015; S. M. Kim et al., 2009; Koles and Budnik, 2012; Osses and Henriquez, 2014; Sanyal et al., 2002; 2003; Sulkowski et al., 2014; Walker et al., 2013). Here we demonstrated a requirement for ROS for activity-dependent structural plasticity at the *Drosophila* NMJ and motoneuron dendrites. Like BMP (Berke et al., 2013) and AP-1 (Milton et al., 2011; Sanyal et al., 2002) ROS have the potential to act as gatekeepers of neuronal plasticity. If acting as a parallel stimulus, ROS signaling at the synapse could be synergistic with other plasticity pathways, which could help overcome issues with biological noise and increase the fidelity of cellular decision making. Alternatively, ROS might deliver requisite post-translational redox modifications to components of other signaling pathways. In support of the latter, ROS regulate by redox modification the activity of the immediate early genes Jun and Fos, which are required for LTP in vertebrates and in *Drosophila* for activity-dependent plasticity of motoneurons, both at the NMJ and central dendrites (Hartwig et al., 2008; Jindra et al., 2004; Loebrich and Nedivi, 2009; Milton et al., 2011; Milton and Sweeney, 2012; Sanyal et al., 2002). ROS modulation of BMP signaling has recently been shown in cultured sympathetic neurons (Chandrasekaran et al., 2015) and of Wnt pathways in non-neuronal cells (Funato et al., 2006; Love et al., 2013; Rharass et al., 2014).

Although we focused exclusively on structural plasticity in the *Drosophila* larva, current evidence suggests synaptic ROS signaling is a conserved and central feature of communication in the nervous system. Several studies have demonstrated a requirement for ROS for Long Term Potentiation (Huddleston et al., 2008; Kamsler and Segal, 2003b; 2003a; Klann, 1998; Knapp and Klann, 2002; Lee et al., 2010) and also correlated defects in learning with altered ROS signaling (Kishida et al., 2006). The underlying cellular and molecular mechanisms though are far from clear, probably because of the complexity of ROS signaling and because of the large number of potential redox sensitive targets. For example, components of the presynaptic release machinery, notably SNAP25, are thought to be modulated by ROS (Giniatullin et al., 2006). Here, we identified DJ-1ß as a ROS sensor in neurons whose interactions with the PTEN/PI3Kinase synaptic growth pathway are governed by redox regulation. This represents a novel mechanism by which ROS, potentially generated as metabolic byproducts, are used to tune connectivity and communication within the nervous system through modifications of neuronal structure. Since ROS can diffuse through membranes these signals have the potential to act non-autonomously, possibly across synaptic partners. Our findings shine a new light on ROS which, when dysregulated with age or under neurodegenerative conditions, might interfere with neuronal adaptive adjustments and thus contribute to network malfunction and synapse loss.

## Competing interests

The authors declare that no competing interests exist.

## Author Contributions

M.O., M.F.Z., S.T.S. & M.L. devised the project. M.O. performed most experiments and analyses. P.B., A.M., M.F.Z. & K.M. generated specimens and dendrite data on activity dependent dendrite growth. A.M. carried out experiments and analysis involving ROS measurements. R.J.H.W. processed specimens for TEM imaging and analysed TEM data. M.O. & M.L. co-wrote the manuscript and S.T.S. & M.L. co-supervised the project.

## Acknowledgements

We would like to thank Nancy Bonini, Tobias Dick, Jörg Grosshans, Karen Hibbard, Fanis Missirlis, Barret Pfeiffer and Alex Whitworth for generous fly stock and reagent donations, Other fly stocks were obtained from the Bloomington Stock Center. The monoclonal antibodies nc82 (anti-Bruchpilot), developed by E. Buchner and colleagues, was obtained from the Developmental Studies Hybridoma Bank, created by the NICHD of the NIH and maintained at The University of Iowa, Department of Biology. We thank Akinao Nose and Hiroshi Kohsaka for kindly sharing figure graphics, Benjamin Risse for his generous support with setting up a FIM Tracker and Matthew Wayland for his generous support with data analysis and imaging. We would also like to thank Jimena Berni, Alex Whitworth and members of the Landgraf lab for valuable comments on the manuscript. This work was supported by BBSRC research grants (BB/IO1179X/1, BB/M002934/1) to ML and (BB/I012273/1, BB/M002322/1) to STS. This work benefited from facilities supported by a Wellcome Trust Equipment Grant (WT079204) and contributions by the Sir Isaac Newton Trust in Cambridge.

## Materials and Methods

### Fly Strains and Husbandry

Wild-type and transgenic strains were maintained on standard yeast–agar–cornmeal medium at 25 °C. The following fly strains were used: *OregonR* and *PTEN*^*CO76*^ (Bloomington Stock Center, Indiana University), *UAS-dTrpA1* (Hamada et al., 2008), *UAS-SOD2* (Missirlis et al., 2003), *UAS-Catalase* (Missirlis et al., 2001), *UAS-Duox* (Ha, 2005), *DJ-1ß*^*Δ93*^ (Meulener et al., 2005), *UAS-DJ-1ß*^*C104A*^ (Meulener et al., 2006), UAS-RNAi lines targeting *SOD1*, *SOD2*, *Catalase* and *PTEN* (KK collection, Vienna Drosophila Resource Centre) (Dietzl et al., 2007), *UAS-PI3K*^*DN*^ (Leevers et al., 1996), *UAS-PTEN* (Gao et al., 2000). The following two GAL4 expression lines were used to target GAL4 to the aCC and RP_2_ motoneurons: *aCC-FLP-GAL_4_* (e*veRN_2_-Flippase, UAS-myr∷mRFP1, UAS-Flp, tubulin8_4_B-FRT-CD_2_-FRT-GAL_4_*) (Roy et al., 2007) expresses GAL4 stochastically in single aCC and RP_2_ motoneurons allowing the imaging of aCC neurons in isolation, as required for dendritic arbor resolution and reconstruction. *aCC/RP_2_-GAL_4_* (*eveRN_2_-GAL_4_* (Fujioka et al., 2003)*, UAS-myr-mRFP_1_, UAS-Flp, tubulin8_4_B-FRT-CD_2_-FRT-GAL_4_; RRaGAL_4_, 20xUAS-6XmCherry∷HA* (Shearin et al., 2014)) was used for NMJ analysis as it expresses GAL_4_ in every aCC and RP_2_ motoneuron. *eveRN_2_-GAL_4_* expression is restricted to the embryo and FLPase-gated *tubulin8_4_B-FRT-CD_2_-FRT-GAL_4_* maintains GAL4 expression at high levels during larval stages.

## Dissection and Immunocytochemistry

### 1^st^ Instar Ventral Nerve Cord (VNC)

Flies were allowed to lay eggs on apple juice-based 518 agar medium for 24 hrs at 25°C. Embryos were dechorionated using bleach (3.5 minutes room temperature) then incubated (25°C) in pre-warmed Sorensen’s saline (pH 7.2, 0.075 M) whilst adhered to a petri dish. Hatched larvae (floating) were recovered hourly and transferred to pre-warmed apple-juice agar plates supplemented with yeast paste. Larvae were allowed to develop for a further 24 hrs (24 hrs after larval hatching, ALH) at 25°C, 27°C or 29°C prior to dissection in Sorensen’s saline. A fine hypodermic needle (30 1/2 G; Microlance) was used as a scalpel to cut off the anterior end of each larva, allowing gut, fat body, and trachea to be removed. The ventral nerve chord and brain lobes, extruded with viscera upon decapitation, were dissected out and transferred to a cover glass coated with poly-L-lysine (Sigma-Aldrich), positioned dorsal side up in Sorensen’s saline. A clean cover glass was placed on top of the preparation, with strips of double-sided sticky tape as spacers positioned along the edges.

### Wandering 3^rd^ Instar

Flies were allowed to lay eggs on apple-juice agar based medium overnight at 25°C, larvae were then incubated at 25°C or 27°C until the late wandering 3^rd^ instar stage. Larvae were reared on yeast paste colored with Bromophenol Blue sodium salt (Sigma-Aldrich) to allow visualization of gut-clearance, an indicator of the late wandering 3^rd^ instar stage. For Di-ethylmaleate (DEM) (Sigma-Aldrich) and paraquat (Sigma-Aldrich) feeding, yeast paste was made using a 5 mM – 15 mM aqueous solution. Larvae were ‘fillet’ dissected in Sorensen’s saline and fixed for 15 minutes at room temperature in 4% formaldehyde (in Sorenson’s saline). Specimens were then washed and stained in Sorensen’s saline containing 0.3% Triton X-100 (Sigma-Aldrich) using the following primary and secondary antibodies; Goat-anti-HRP Alexa Fluor 488 (1:400) or Goat-anti-HRP Alexa Fluor 594 (1:400) (Jackson ImmunoResearch Cat. Nos. 123-455-021 or 123-585-021), Rabbit-anti-DsRed (1:1200) (ClonTech Cat. No. 632496), Donkey-anti-Rabbit CF568 (1:1200) (Biotium Cat. No. 20098), Mouse-nc82 (1:100) (Developmental Studies Hybridoma Bank), Donkey-anti-Mouse Alex Fluor 488 (1:600) (ThermoFisher Cat. No. A-21202) or Donkey-anti-Mouse CF633 (1:300) (Biotium Cat. No. 20124) incubated overnight at 4°C or 2 hrs at room temperature. Specimens were mounted in EverBrite mounting medium (Biotium Cat. No. 23001).

## Image Acquisition and Analysis

### 1^st^ Instar Ventral Nerve Cord (VNC)

Ventral nerve cords were pre-screened for fluorescently labeled, isolated, aCC motoneurons using a Zeiss Axiophot compound epifluorescence microscope and a Zeiss Plan-Neofluar 40x/0.75 N.A. objective lens. Suitable VNCs were imaged immediately with a Yokagawa CSU-22 spinning disk confocal field scanner mounted on an Olympus BX51WI microscope, using a 60×/1.2 N.A. Olympus water immersion objective. Images were acquired with a voxel size of 0.2 × 0.2 × 0.3 µm using a QuantEM cooled EMCCD camera (Photometrics), operated via MetaMorph software (Molecular Devices). Dendritic trees were digitally reconstructed using Amira Resolve RT 4.1 (FEI), supplemented with a 3D reconstruction algorithm (Evers et al., 2005; Schmitt et al., 2004), and images were processed using Amira (FEI) and ImageJ (National Institutes of Health).

### Wandering 3^rd^ Instar

Dissected specimens were imaged using a Leica SP5 point-scanning confocal, and a 63x/1.3 N.A. (Leica) glycerol immersion objective lens and LAS AF (Leica Application Suite Advanced Fluorescence) software. Confocal images were processed using ImageJ and Photoshop (Adobe). Bouton number of the NMJ on muscle DA1 [1] from segments A3-A5 was determined by counting every distinct spherical varicosity along the NMJ branch. DA1 muscles were imaged using a Zeiss Axiophot compound microscope and a Zeiss Plan-Neofluar 10x/0.3 N.A. objective lens. Muscle surface area (MSA) was determined by multiplying muscle length by width using ImageJ. In order to correct for subtle differences in animal size (typically 5-10%) bouton number normalization was performed using the following calculation: (mean control MSA / mean experimental MSA) x test bouton number = normalized experimental bouton number.

### Ratiometric ROS Reporter

*aCC/RP2-Gal4* was used to drive the expression of *UAS-mito-roGFP2-Orp1* (Albrecht et al., 2011; Gutscher et al., 2009) in all aCC and RP2 motoneurons. Wandering third instar larvae were fillet dissected in PBS-NEM (137mM NaCl, 2.7mM KCl, 10mM Na2HPO4, 1.8mM KH2PO4, 20mM N-ethylmaleimide (NEM), pH 7.4). Larval fillet preparations were incubated for 5 minutes in PBS-NEM then fixed for 8 minutes in 4% formaldehyde (in PBS-NEM). Specimens were washed three times in PBS-NEM and then equilibrated in 70% glycerol. Specimens were mounted in glycerol and imaged the same day. Imaging was performed on a Leica SP5 point-scanning confocal, using a 63x/1.3 N.A. (Leica) glycerol immersion objective lens. The reporter was excited sequentially at 405nm and 488nm (Albrecht et al., 2011) with emission detected at 500– 535nm. 16-bit images were acquired using Leica LAS AF software and processed using ImageJ. Z-stack images were maximally projected and converted to 32-bit. To remove fringing artefacts around bouton edges 488nm images were thresholded using the ‘Intermodes’ algorithm with values below threshold set to ‘not a number’, and ratio images were created by dividing the 405nm image by the 488nm image pixel by pixel (Albrecht et al., 2011). Regions of Interest were taken on the ratio image spanning the entire NMJ and the mean value obtained from each NMJ was used for statistical analysis.

### Transmission Electron Microscopy

Third instar wandering larvae were fillet dissected in PBS and fixed overnight in 0.1M NaPO_4_, pH 7.4, 1% glutaraldehyde, and 4% formaldehyde, pH 7.3. Fixed specimens were washed 3× in 0.1M NaPO_4_ before incubation in OsO_4_ (1% in 0.1M NaPO4; 2hr). Preparations were washed 3× in distilled water, incubated in 1% uranyl acetate, then washed again 3× in distilled water and dehydrated through a graded ethanol series: 20% increments starting at 30% followed by two 100% changes and then 2× 100% propylene oxide. Specimens were incubated in a graded series of epon araldite resin (in propylene oxide): 25% increments culminating in 3× 100% changes. Individual muscles were then dissected and transferred into embedding molds, followed by polymerization at 60°C for 48 hrs. Resin mounted specimens were sectioned (60–70 nm) using glass knives upon a microtome (Ultracut UCT; Leica). Sections were placed onto grids, incubated in uranyl acetate (50% in ethanol), washed in distilled water and incubated in lead citrate. Sections were imaged using a transmission electron microscope (TECNAI 12 G^2^; FEI) with a camera (Soft Imaging Solutions MegaView; Olympus) and Tecnai user interface v2.1.8 and analySIS v3.2 (Soft Imaging Systems).

### Behavior – larval crawling analysis

To record larval crawling, mid-3^rd^ instar larvae (72hrs ALH) were briefly rinsed in water to remove any food and yeast residues, then up to 12 larvae were placed into a 24cm x 24cm arena of 0.8% agar in water, poured to 5 mm thickness. Crawling behavior was recorded in a temperature and humidity controlled incubator at temperatures ranging from 25-32^o^C, as indicated for each experiment. Larvae were allowed to acclimatise for 5 minutes, then recorded for 15 minutes under infrared LED illumination (intensity from 14.33 nW/mm^2^ in the edge to 9.12 nW/mm^2^ in the center), using frustrated total internal reflection using a modified FIM tracker (Milton et al., 2011; Risse et al., 2013) https://www.uni-muenster.de/PRIA/en/FIM/index.html. Larvae were recorded with a Basler acA2040-180km CMOS camera using Pylon and StreamPix software, mounted with a 16mm KOWA IJM3sHC.SW VIS-NIR Lens. Images were acquired at 5 frames per second. For each larvae, average crawling speed was calculated from long, uninterrupted forward crawls identified manually using FIMTrack. The 15 minute recording period was partitioned into 5 minute sections with each larvae being assayed once within each section, allowing each specimen to be sampled a maximum of 3 times. We observed no change in average crawling speed within the duration of the 15-minute recording.

#### Data Analysis

All data handling was performed using Prism software (GraphPad). NMJ bouton number and ratiometric ROS reporter data were tested for normal / Gaussian distribution using the D’Agostino-Pearson omnibus normality test. Due to a lower replicate number, dendritic arbor reconstruction data were tested for normality using the Kolmogorov-Smirnov with Dallal-Wilkinson-Lilliefor P value test. Normal distribution was thus confirmed for all data presented, which were compared using one-way analysis of variance (ANOVA), with Tukey’s multiple comparisons test where *P<0.05, **P<0.01, ***P<0.001, ****P<0.0001.

---

**Figure 1-Figure Supplement 1:**
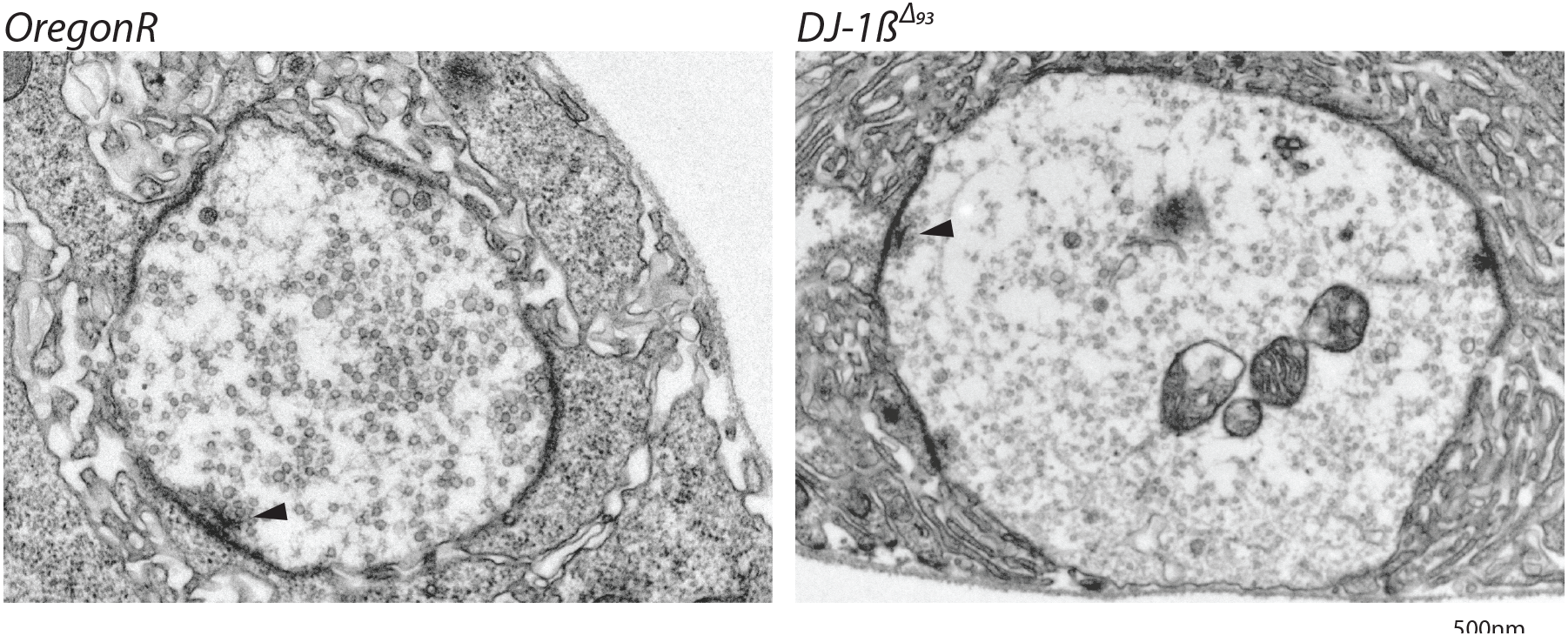
Prolonged over-activation targeted to motoneurons (*D42-GAL4*; *UAS-dTrpA1*) also leads to adaptation with reduced crawling speed (dTRPA1 channels open at 27°C, closed at 22°C). Control is *D42-GAL4 / +.* Each data point represents the crawling speed from an individual uninterrupted continuous forward crawl, n = specimen replicate number, up to 3 crawls measured for each larva. ****P<0.0001 students t test.

**Figure 4-Figure Supplement 1:**
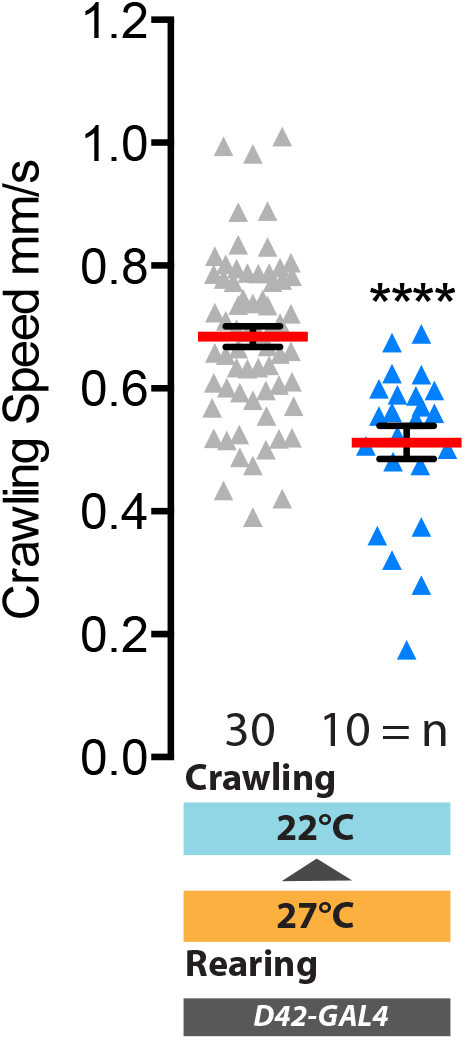
*DJ-1β*^*Δ93*^ mutant 3^rd^ instar larval NMJs are phenotypically normal. Representative TEM bouton cross-sectional images showing that pre-synaptic architecture is intact including active-zones with associated clustered synaptic vesicles (arrowed).

**Figure 4-Figure Supplement 2:**
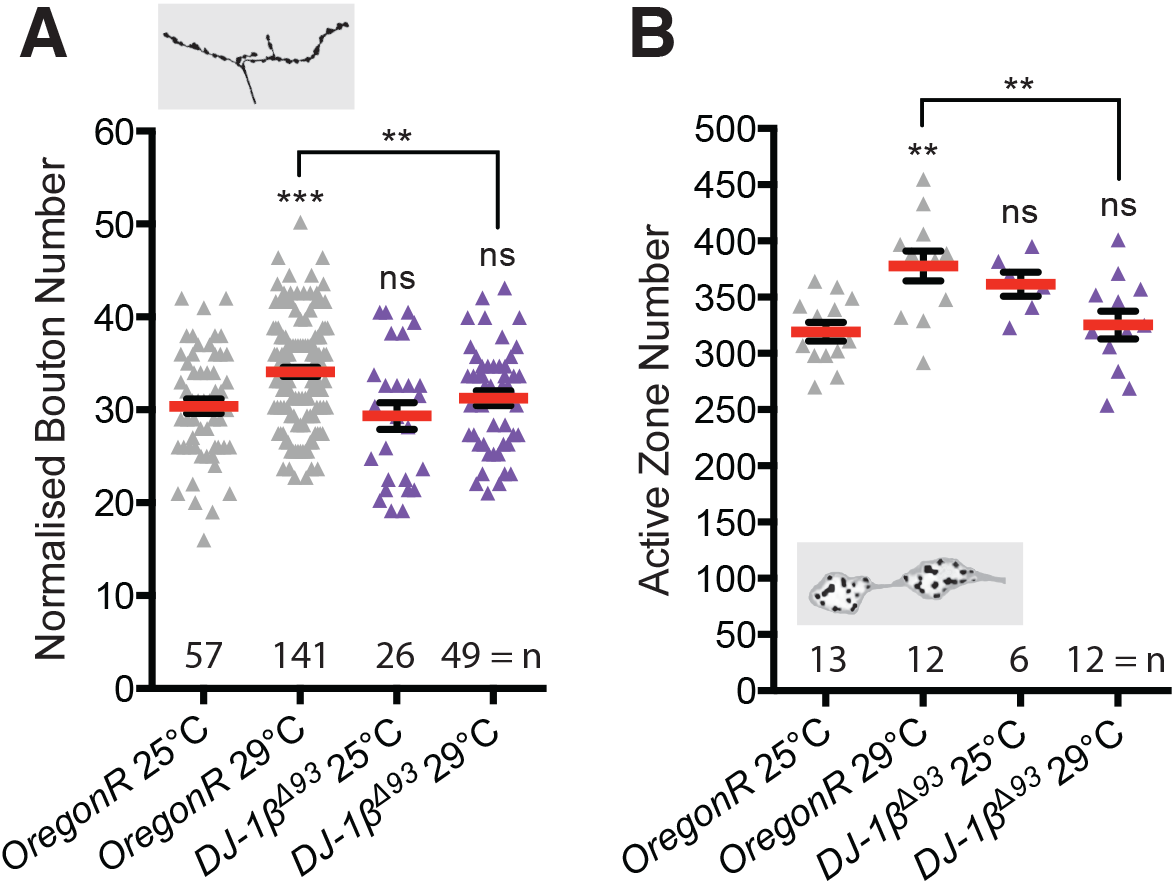
**A**. *DJ-1β* null mutant (*DJ-1β*^*Δ93*^) larvae do not show systemic activity-dependent NMJ elaboration. Normalised bouton number dot plot showing significantly increased NMJ elaboration in *OregonR*, but not *DJ-1β*^*Δ93*^, larvae reared at 29°C vs 25°C. **B**. *DJ-1β*^*Δ93*^ larvae also do not show systemic activity-dependent increase in active zone number. *OregonR*, but not *DJ-1β*^*Δ93*^, show significantly increased NMJ active zone number when reared at 29°C vs 25°C. Mean +/-SEM, ANOVA **p<0.01, ***p<0.001, n=replicate number.

